# The small mycobacterial ribosomal protein, bS22, modulates aminoglycoside accessibility to its 16S rRNA helix-44 binding site

**DOI:** 10.1101/2023.03.31.535098

**Authors:** Soneya Majumdar, Ayush Deep, Manjuli R. Sharma, Jill Canestrari, Melissa Stone, Carol Smith, Ravi K. Koripella, Pooja Keshavan, Nilesh K. Banavali, Joseph T. Wade, Todd A. Gray, Keith M. Derbyshire, Rajendra K. Agrawal

## Abstract

Treatment of tuberculosis continues to be challenging due to the widespread latent form of the disease and the emergence of antibiotic-resistant strains of the pathogen, *Mycobacterium tuberculosis*. Bacterial ribosomes are a common and effective target for antibiotics. Several second line anti-tuberculosis drugs, e.g. kanamycin, amikacin, and capreomycin, target ribosomal RNA to inhibit protein synthesis. However, *M. tuberculosis* can acquire resistance to these drugs, emphasizing the need to identify new drug targets. Previous cryo-EM structures of the *M. tuberculosis* and *M. smegmatis* ribosomes identified two novel ribosomal proteins, bS22 and bL37, in the vicinity of two crucial drug-binding sites: the mRNA-decoding center on the small (30S), and the peptidyl-transferase center on the large (50S) ribosomal subunits, respectively. The functional significance of these two small proteins is unknown. In this study, we observe that an *M. smegmatis* strain lacking the *bs22* gene shows enhanced susceptibility to kanamycin compared to the wild-type strain. Cryo-EM structures of the ribosomes lacking bS22 in the presence and absence of kanamycin suggest a direct role of bS22 in modulating the 16S rRNA kanamycin-binding site. Our structures suggest that amino-acid residue Lys-16 of bS22 interacts directly with the phosphate backbone of helix 44 of 16S rRNA to influence the micro-configuration of the kanamycin-binding pocket. Our analysis shows that similar interactions occur between eukaryotic homologues of bS22, and their corresponding rRNAs, pointing to a common mechanism of aminoglycoside resistance in higher organisms.

## Introduction

Tuberculosis (TB) was the leading cause of death due to infectious diseases until the emergence of SARS-CoV-2 (1). The current treatment regimen for TB requires patients to take a multi-antibiotic cocktail for at least six months (1). This extended treatment program often results in patient non-compliance and the subsequent emergence of drug-resistant *Mycobacterium tuberculosis* (2). Several first- and second-line anti-tubercular drugs are ineffective against these resistant strains of *M. tuberculosis* (3). Aminoglycoside (AG) antibiotics (kanamycin, amikacin, and capreomycin) have been extensively used as second-line injectables for treating multidrug-resistant tuberculosis (MDR-TB) for decades (4). These are polycationic, pseudo-oligosaccharides with a neamine core (5). The most clinically used and investigated AGs have a neamine core of 2-deoxystreptamine (2-DOS or ring-II (RII)) linked to amino sugars through glycosidic linkages (5). The AG subclasses have different chemical linkages or substitutions on RII (**Fig. 1a**). For example, the kanamycin subclass, which includes dibekacin, amikacin, arbekacin, and tobramycin, are 4,6-disubstituted 2-deoxystreptamines (6) having ring-I (RI) and ring-III (RIII) attached at positions 4 and 6 of 2-DOS/RII, respectively (**Fig. 1a**). These drugs are broad-spectrum antibiotics that inhibit bacterial protein synthesis and have bactericidal properties (6).

**Figure 1.**
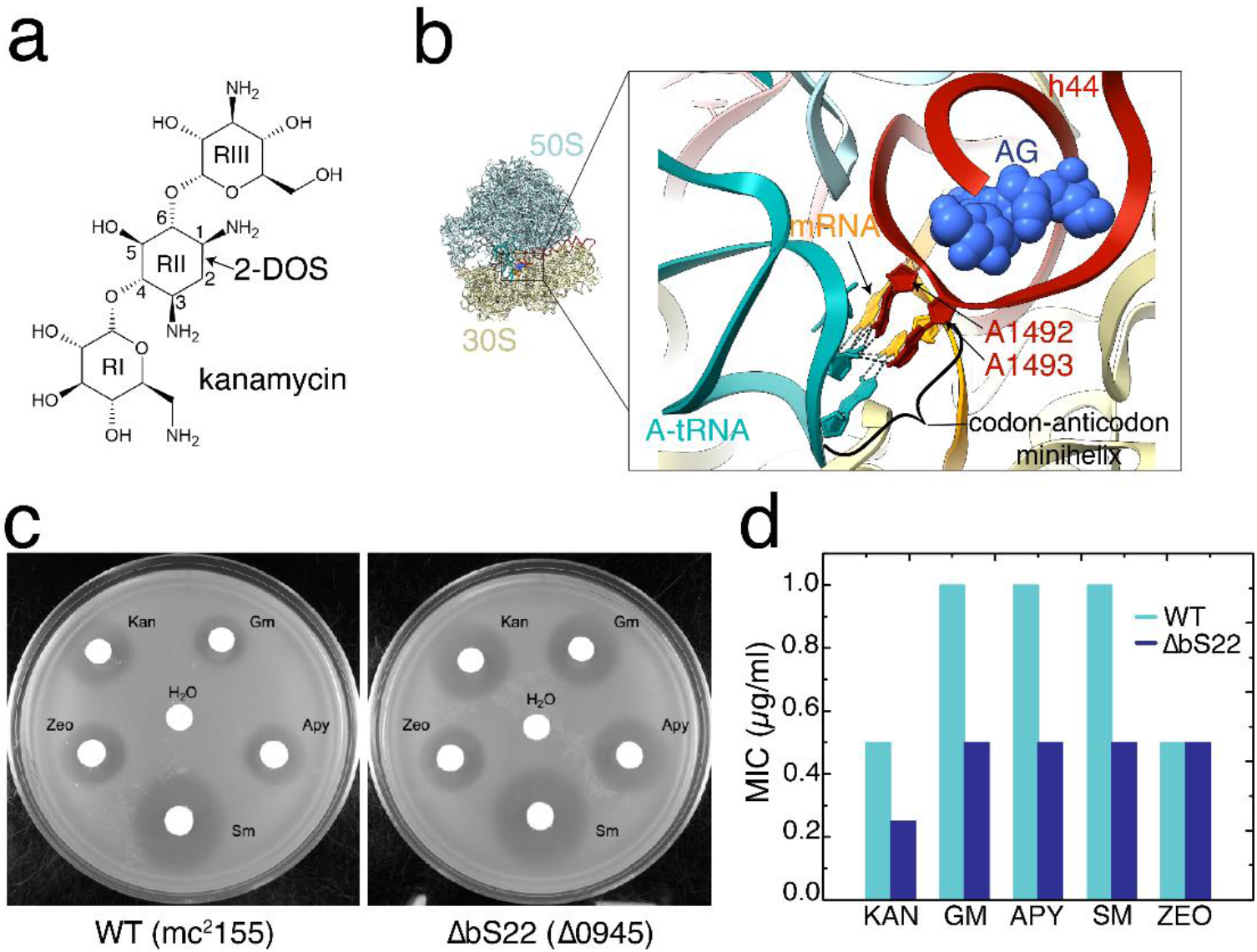
Effect of aminoglycosides on ΔbS22 ribosomes. (a) kanamycin — a 4,5-linked 2-deoxystreptamine (2-DOS) aminoglycoside. (b) Overview of the aminoglycoside-binding pocket in the small ribosomal subunit’s 16S rRNA helix 44 (h44, maroon). Binding of aminoglycosides (AG, blue) flips the monitoring bases (*E. coli* numbers A1492 and A1493, maroon). The flipped monitoring bases interact with the minihelix formed between codon (teal) and anticodon (mRNA, gold) nucleotides. (c) Disk-diffusion assay showing increased sensitivity to aminoglycosides of a Δ*bS22* strain of *M. smegmatis* in contrast to the wild-type strain, mc^2^155. For each antibiotic, equivalent amounts were soaked into filter disks placed on a lawn of cells in top agar. The zeocin (control) shows sensitivity specific to aminoglycosides, which bind to the 30S subunit of the ribosome. A *bL37* deletion had no effect on sensitivity to these antibiotics (data not shown). (d) MIC values comparing the sensitivity to aminoglycosides, kanamycin (KAN), gentamicin (GM), apramycin (APY), and streptomycin (SM) of a Δ*bs22* strain (ΔbS22) of *M. smegmatis* in contrast to the wild-type (WT) strain, mc^2^155. Zeocin (ZEO), a bleomycin derivative with a DNA-binding mode of action, is used as a control.

AGs bind primarily to helix 44 (h44) of the 16S ribosomal RNA (rRNA) near the aminoacyl-tRNA binding site (A-site) (7) within the small (30S) subunit (SSU) (**Fig. 1b**). Point mutations in h44 at nucleotide residues (nts) A1408, G1491, U1495 (8-10), or methylation at nts G1405 or A1408 (11) are sufficient to confer AG resistance. These mutations affect the interactions stabilizing RI and/or RII of the drug molecule compromising its binding (7). Cryo-electron microscopic (cryo-EM) and X-ray crystallographic structures of various AGs in complex with the ribosome show that AGs with a 2-DOS ring bind h44 in a similar manner (12-15). AG binding occludes the two codon-anticodon minihelix-monitoring bases, A1492 and A1493, that attain a “flipped out” conformation from the h44 internal loop (**Fig. 1b**) (16). In the absence of AGs, the monitoring bases attain the flipped-out conformation only to interact with the first and second bases of the codon-anticodon minihelix formed between a mRNA codon and its cognate aminoacyl-tRNA (**Fig. 1b**). This step known as proof-reading is crucial for discrimination of cognate versus near-cognate tRNAs at the A-site. AG binding at h44 reduces the energy barrier for conformational switching of the monitoring bases, which stabilizes the A-site tRNA irrespective of whether it is cognate or near-cognate (16). Thus, AG binding affects protein synthesis by; (i) increasing promiscuous incorporation of near-cognate amino-acyl tRNA, and (ii) inhibiting tRNA-mRNA translocation by resisting disruption of the codon-anticodon duplex when domain-IV of EF-G tries to insert itself into the A-site.

Cryo-EM structures of *Mycobacterium smegmatis* and *M. tuberculosis* ribosomes revealed two additional small proteins — bL37 in the large (50S) ribosomal subunit (LSU) and bS22 in the SSU (17-19). bL37, encoded by MSMEG_1916, is a small highly basic (pI=12.02) protein of 24 amino acids and is conserved throughout the *Actinobacteria*. Its *M. tuberculosis* ortholog is not annotated, but it is encoded from H37Rv, genome coordinates 3,597,967-3,598,041 nts. bS22, encoded by MSMEG_0945 (*Rv0550B*), is a highly basic (pI=12.75) protein of 33 aa, which is conserved throughout the actinobacteria, yeast and in eukaryotes (17, 20, 21). bL37 and bS22 lie in the vicinity of the catalytic (peptidyltransferase) and decoding centers, respectively, suggesting roles in protein translation, and effects on anti-tubercular drugs that interact with ribosome catalytic centers (17-19).

Here we have investigated the potential roles of bL37 and bS22 in the ribosome using molecular, genetic and structural approaches. We show that an *M. smegmatis ΔbS22* derivative is more susceptible to kanamycin than wild-type or *ΔbL37* strains. By determining the cryo-EM structures of ΔbS22-*M. smegmatis* ribosomes in the presence and absence of kanamycin, we identified the molecular interactions responsible for kanamycin binding and determined how those interactions are impaired by bS22. In support of this model, we identify a single amino-acid residue in bS22 that is key in modulating resistance to kanamycin.

## Results

### Characterization of *M. smegmatis* strains lacking bL37 and bS22

To begin to address the roles of bL37 and bS22, we made precise deletions of each gene by recombineering (22). Each gene was initially replaced with a cassette encoding zeocin resistance, and this cassette was subsequently excised by the Cre recombinase, leaving a *loxP* scar (23). A strain containing both unmarked-gene deletions was also created. The three deletion strains were then put through a battery of assays to determine the impact of the deletions on cell viability and growth compared with a wild-type control. These assays included standard growth curves in different medium and co-culture growth competition assays with the wild-type strain involving serial dilution and growth over four days before determining relative bacterial counts (Materials and Methods). There was no significant difference in relative growth of the three deletion strains either in mono- or co-culture compared with a differentially marked wild-type strain. These assays were completed in both rich and minimal media and at temperatures ranging from 30^°^C to 45^°^C. In addition, all strains grew mature, comparable biofilms in Sauton’s minimal medium. Mycobacteria encode thousands of leaderless genes; approximately 1/3 of the 5’mRNAs are leaderless (LL) (24-26). LLmRNAs lack a leader sequence with a ribosome-binding site, and instead transcription and translation initiate at the 5’-end of an mRNA beginning RUG (25). Thus, we also examined whether bS22 or bL37 facilitate translation of LLmRNAs, as both are absent in *Escherichia coli* and similar species that also lack LL genes. We tested the effect of deleting MSMEG_1916 or MSMEG_0945 on RNA levels and ribosome-associated RNA levels for all genes in *M. smegmatis* using RNA-seq and Ribo-seq, respectively. We observed no major differences in expression RNA levels or ribosome-associated RNA levels for any group of RNAs, including LLmRNAs (**Fig. S1**). In addition, there were no unique subsets of genes that were differentially expressed in the mutant strains compared to wildtype. In summary, under the laboratory conditions tested, we failed to establish any defect in growth or global translation in strains lacking either bL37 or bS22 or both.

### bS22 reduces sensitivity to aminoglycosides

The mycobacterial ribosomal protein bS22 lies adjacent to the aminoglycoside (AG)-binding pocket within h44 of the 16S rRNA. We therefore hypothesized that our bs22 deletion strain would exhibit altered sensitivity to AG antibiotics. To test this hypothesis, we screened the sensitivity of the *ΔbS22* strain to a variety of ribosome-targeting antibiotics using a zone of inhibition assay including apramycin, azithromycin, clarithromycin, chloramphenicol, gentamycin, kanamycin, retapamulin, streptomycin and tetracycline. The AG antibiotics, apramycin, kanamycin and gentamycin reproducibly resulted in larger zones of inhibition compared to the other antibiotics and controls (**Fig. 1c**). This increased sensitivity was observed with both the *Δbs22* and the double-deletion *ΔbL37ΔbS22* strains, but not with *ΔbL37 M. smegmatis*, indicating there is no synergistic effect between the small proteins conferring resistance to these antibiotics. Below, we have focused on the role of bS22.

To more rigorously confirm this change in antibiotic sensitivity, we determined the minimum inhibitory concentrations (MIC) for five ribosomal targeting antibiotics (apramycin, chloramphenicol, kanamycin, gentamycin and streptomycin) and zeocin (as a non-ribosomal antibiotic) against the *ΔbS22* strain. The mutant strain was 2-fold more sensitive to apramycin, kanamycin, gentamycin and streptomycin than the wild-type control (**Fig. 1d**). Sensitivity to both chloramphenicol and zeocin was unchanged in the strain lacking the bS22 protein, similar to the observations made with the disk diffusion assays.

### Structure of kanamycin bound ΔbS22 *M. smegmatis* small ribosomal subunit

We determined the cryo-EM structures of the wild-type and ΔbS22 *M. smegmatis* small (30S) ribosomal subunit (SSU) in the presence and absence of kanamycin to a global resolution of 3.1 Å (**Fig. S2-S4**). In the cryo-EM maps of both complexes, we observed well-defined densities for the SSU body, including h44 and bS22 (for wildtype ribosomes, **Fig. S4**), while the head domain was relatively dynamic. We could model the entire 16S rRNA and most ribosomal S-proteins, except bS1 and bS22 in the ΔbS22 mutant ribosome. Representative segments displaying high-resolution features of either map are shown (**Fig. S5**). Loss of bS22 does not affect the overall architecture of the SSU. However, in the absence of bS22 there were subtle structural changes in the sugar-phosphate backbone of the neighboring 16S rRNA helices h27, h44, and h45.

The cryo-EM map of the ΔbS22 SSU-kanamycin complex allows the unambiguous identification of kanamycin at the ribosomal A-site within 16S rRNA (**Fig. 2ai**). Kanamycin binds to the base of h44 and stabilizes a flipped-out conformation of the two, universally conserved, monitoring rRNA nts, A1476 and A1477 (A1492 and A1493, respectively, in *E. coli*) (**Fig. 2ai**). In contrast, the cryo-EM map of ΔbS22 SSU lacking kanamycin, shows that A1476 and A1477 are distinctly flipped-in (**Fig. 2aii**). This 16S rRNA site corresponds to the known deoxystreptamine aminoglycoside-binding pocket in bacterial ribosomes (12-15). The three-ring structures of kanamycin each formed clear stabilizing interactions within the conserved binding pocket. The RI ring stacks with nt G1475, which appropriately positions its 6’-NH2 to interact with nts A1391 and C1392 (**Fig. 2b**). The RII N1 and N5 amino groups of kanamycin interact with nts G1478 and U1479, respectively, and the O5 hydroxyl with nts G1478 and A1391 (**Fig. 2b**). The RIII ring of kanamycin establishes multiple interactions; it forms hydrogen bonds with N7 and O6 (Hoogsteen sites) of nt G1388 and with the sugar-phosphate backbone, O4 and N4 of the nts U1389, U1479, and C1480, respectively (**Fig. 2b**).

**Figure 2.**
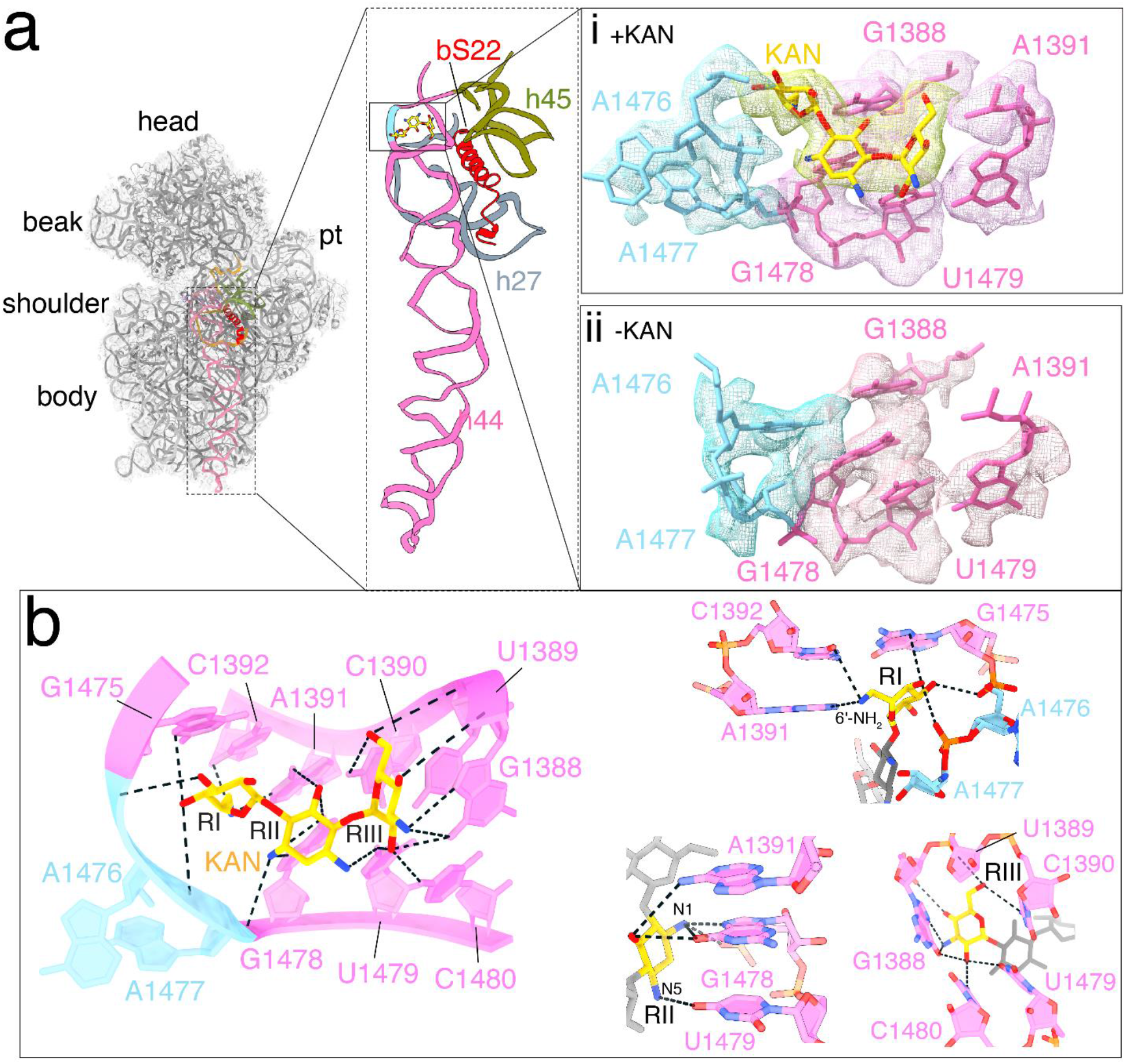
Cryo-EM structure of the ΔbS22 *M. smegmatis* small ribosomal subunit in the presence and absence of kanamycin. Cryo-EM density of the aminoglycoside-binding pocket in the (i) presence and (ii) absence of kanamycin (yellow). The monitoring bases A1476 and A1477 (pink) are flipped out upon drug binding (i) compared to (ii). (c) left-Interactions of the ribosomal A-site (blue) components with kanamycin, right-Interactions stabilizing the individual rings (RI-RIII) of kanamycin.

### bS22 alters the aminoglycoside binding pocket in h44

To understand the molecular basis for aminoglycoside sensitivity, we compared the SSU structures from the wild-type and *ΔbS22* strains in the presence and absence of kanamycin (**Fig. 3a**). The structure of the wild-type ribosome suggests that salt bridges between key lysine residues of bS22 and interacting rRNA phosphate groups stabilize bS22 binding. A salt bridge between Lys16 from bS22 and the sugar-phosphate backbone of h44 at nt U1389, amid the cluster of AG interacting nts 1388-1392, further suggests a structural basis for the functional consequence of bS22 binding. Comparison with the ΔbS22 ribosome showed that the absence of bS22 alters the conformation of the kanamycin-interacting nts, 1388 and 1389 (**Fig 3b-f**, compare gray with sea green and pink). In the ΔbS22 ribosome, G1388 lies in a different plane, ∼1.4 Å from its wild-type location. In the absence of bS22, the sugar-phosphate backbone across nts 1388-1392 shifts toward the kanamycin-binding pocket (**Fig. 3a**). Further, in the absence of bS22, U1389 is rotated by 8.7°, deflecting the base 2.5 Å away from bS22 toward the kanamycin-binding pocket (**Fig. 3e**). The observed conformational changes in the ΔbS22 ribosome of the nts 1388-1392 (which interact directly with kanamycin, **Fig. 2b**) moves these nts towards the AG-binding pocket and could stabilize kanamycin binding. Our collective structural data show that a consequence of bS22 binding is conformational alteration of the kanamycin-binding pocket, consistent with the increased AG sensitivity observed by ΔbS22 strains (**Fig 1d**).

**Figure 3.**
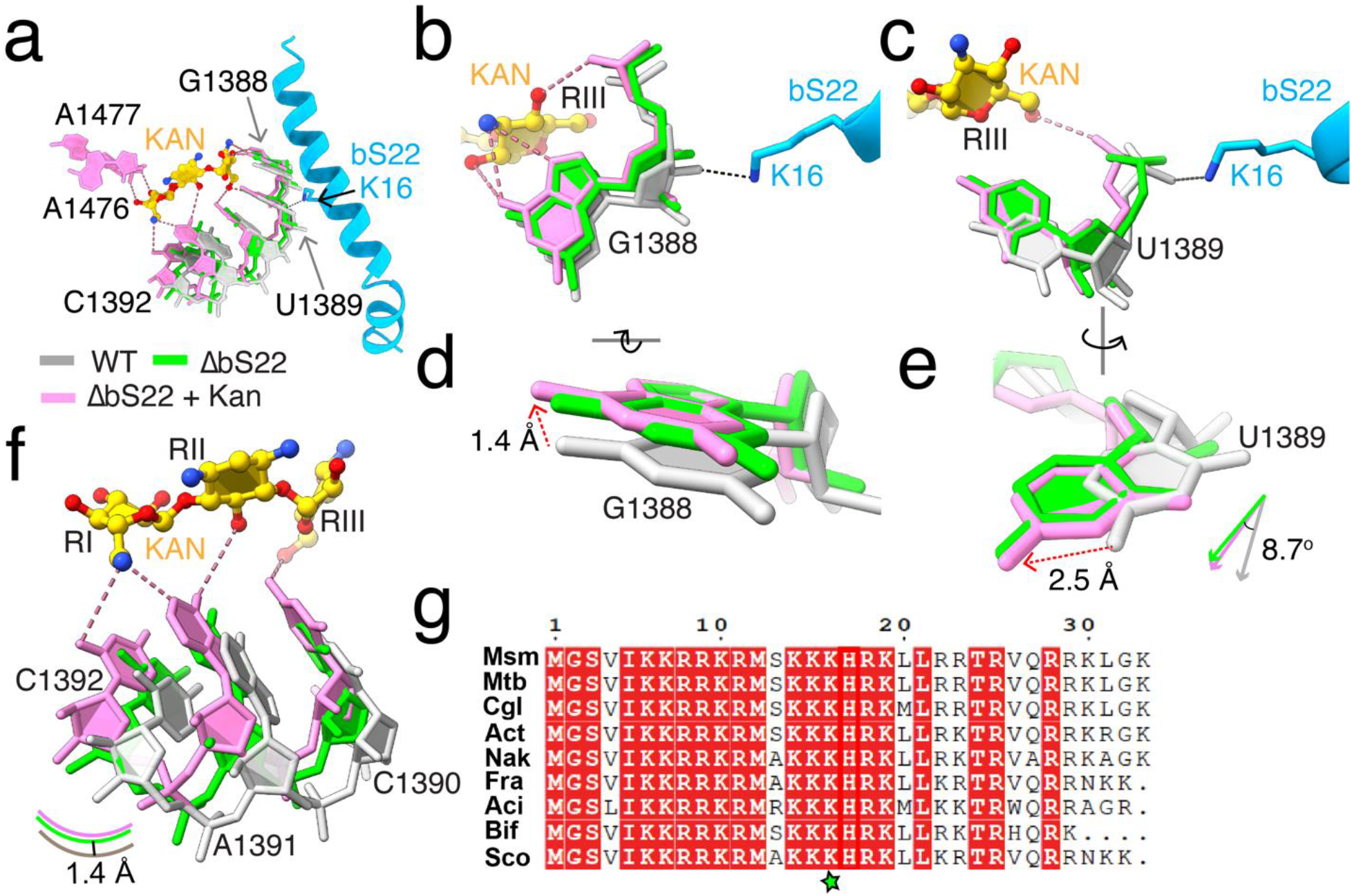
Interactions of *M. smegmatis* bS22 with h44. (**a**) Superimposition of structures of the *M. smegmatis* kanamycin-binding pocket extracted from the wild-type (grey) and ΔbS22 small ribosomal subunits in the presence (pink) and absence of kanamycin (sea green). Interaction of bS22-Lys16 with the sugar-phosphate backbone of G1388-U1389 is highlighted. The conformational changes of the kanamycin interacting nucleotides (b) G1388 (rotated and zoomed in d), (c) U1389 (rotated and zoomed in e) and, (f) the C1390-C1392 segment between wild-type and ΔbS22 ribosomes are shown. **(g)** Multiple sequence alignment of bS22 homologs from *M. smegmatis* (Msm), *M. tuberculosis* (Mtb), *Corynebacterium glutamicum* (Cgl), *Actinosynnema* (Act), *Nakamurella* (Nak), *Frankia* (Fra), *Acidimicrobium* (Aci), *Bifidobacterium* (Bif), and *Streptomyces coelicolor* (Sco). The Lys16 (K16) residue which interacts with U1389 is indicated by a green star.

bS22, including its amino-acid residue Lys16, which interacts with kan-binding pocket in h44, is highly conserved across mycobacteria and distant Actinobacteria (**Fig. 3g**). Notably, conservation of bS22 is also observed in the N-terminal region of the yeast Cox24p, the mammalian mitochondrial protein mS38, and the eukaryotic ribosomal protein eL41 (27-31). Superimposition of mitochondrial mS38 (31), *S. cerevisiae* Cox24p (20) ribosomal protein eL41 and *M. smegmatis* SSUs structures further establishes the overall conservation. We note that Lys142 from mS38 and Arg9 from eL41 interact with their corresponding rRNA residues within h44 similar to that of Lys16 of bS22 (**Fig. 4a, b**). This broad consensus suggests an important functional role for the stabilizing effects conferred by a salt bridge in this ribosomal context.

**Figure 4.**
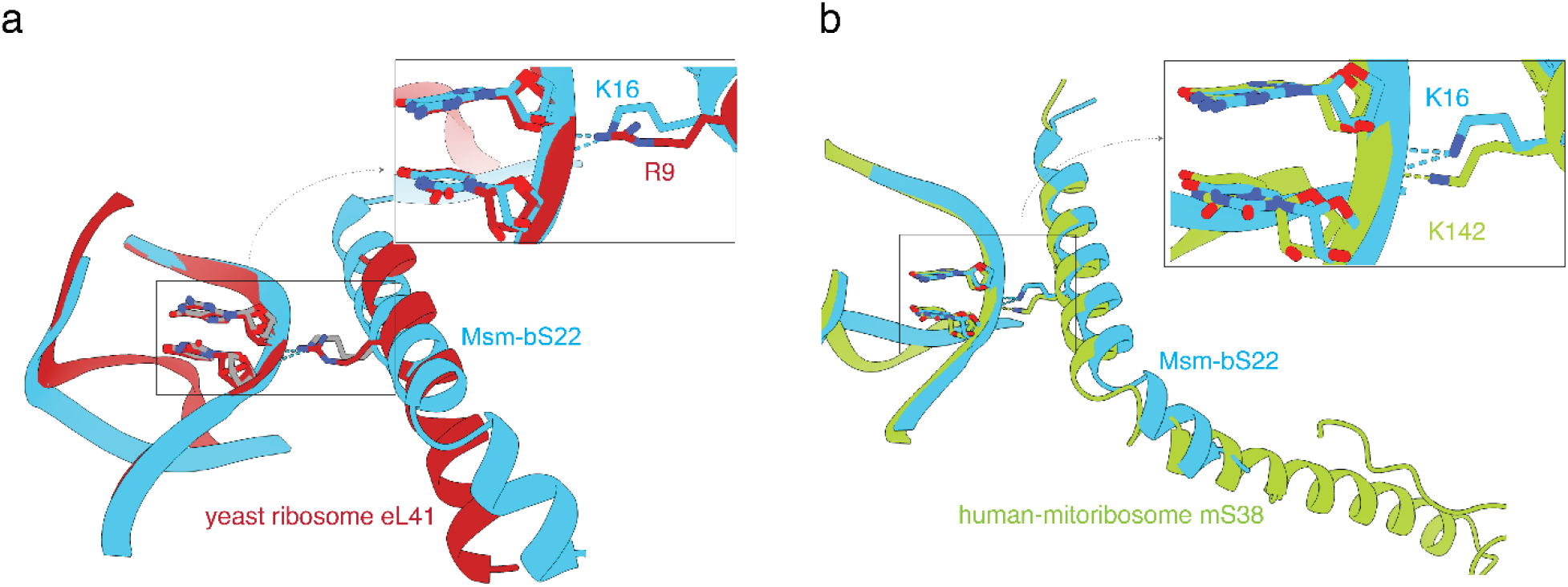
Interactions of bS22 homologous proteins eL41 and mS38 of cytoplasmic and mitochondrial ribosomes with h44. **(a)** Superimposition of the rRNA and bS22 homolog regions from the small ribosomal subunit structures of *M. smegmatis* (cyan) and yeast cytoplasmic ribosome (red, PDB:4V88). eL41-R9 is positioned and interacts with h44 similarly to S22-K16. (**b**) Superimposition of the rRNA and S22 homolog regions from the small ribosomal subunit structures of *M. smegmatis* (cyan) and human mitoribosome (green, PDB:7L08). Interaction of mS38-K142 with h44 mimics S22-K16-h44 interaction (inset).

## Discussion

The Mycobacterial ribosome-specific protein bS22 lies near the decoding center in the SSU similarly placed to its orthologs (mS38, Cox24p and eL41) in the mitochondrial and eukaryotic ribosomes (17-19). Eukaryotic orthologs of bS22 are often significantly larger, varying in size from 111 to greater than 500 amino acids, but they all include the conserved DUF1713 domain at the C-terminus that is present in bS22 (https://www.ncbi.nlm.nih.gov/Structure/cdd/PF08213). This highly conserved domain is rich in lysines and arginines, which ribosome structures show interacting with multiple, neighboring rRNA helices, including the rRNA helix 44 that forms the decoding site. These proteins also form inter-subunit bridges with the ribosomal large subunit (LSU).

Bacteria were not thought to encode mS38 orthologs until they were discovered embedded in the mycobacterial ribosome (17-19). In RNA and ribosomal-profiling experiments, we independently detected high levels of gene expression in both *M. smegmatis* and *M. tuberculosis* of the *bs22* gene (MSMEG_0945 and Rv0550B, *respectively*). The conservation of bS22 homologs in Actinobacteria and eukaryotes, combined with its absence in many bacteria including *E. coli*, prompted us to investigate the role of bS22 in the model mycobacterium *M. smegmatis*. We generated a precise deletion of bS22 (MSMEG_0945), establishing that it is non-essential. We also determined that the relative growth rates of the wild-type and *ΔbS22* strains are comparable under both optimal- and competitive-growth conditions. The functionality of the mycobacterial ribosome lacking bS22 was further assessed by performing RNA- and Ribo-footprinting analyses. Comparing profiles of the wild-type and *ΔbS22* strains, showed that there were no significant differences in (i) global gene expression, (ii) relative levels of translation, or (iii), expression of leaderless mRNAs compared with leadered mRNAs. We found no genes that were differentially expressed in the mutant strains compared to wildtype.

The eukaryotic eL41 protein is also not essential; a ΔeL41 yeast strain has no growth defect compared with the wild-type strain. Despite its non-essentiality, eL41 is considered important for subunit association as it is predicted to stabilize the inter-subunit bridges between eB12 and eB13 (32). It was proposed that eL41 interacts with specific mRNAs along with the elements of the mRNA exit channel to initiate translation. Mitochondrial ribosomes lacking mS38 synthesizes all proteins, but at an attenuated rate (33). Mitochondrial mRNAs are often leaderless or lack a Shine-Dalgarno sequence and it was further hypothesized (but not supported by experimental data) that mS38 might specifically facilitate ribosome-binding to these atypical mRNAs (33). Our observations in mycobacteria that show loss of bS22 has no detectable impact on ribosome occupancy of leaderless mRNAs.

bS22 directly interacts with h44 and lies adjacent to the AG-binding pocket, thus, it was speculated that it might modulate AG-binding (17). Indeed, we observe that *M. smegmatis* lacking bS22 is more sensitive to kanamycin (and related AGs) as compared with the wild-type control. We determined the structures of both wild-type SSUs and those lacking bS22, in the presence and absence of kanamycin. These structures allowed us to define a critical h44 interaction with bS22 that impacts kanamycin binding to h44. We found that K16 of bS22 interacts directly with the sugar-phosphate backbone of G1388-U1389. In ribosomes lacking bS22, the h44 nts 1388-1392 lie in a closer proximity to kanamycin compared to the wild-type ribosomes that could promote stable binding. Previous studies in organisms that inherently lack bS22 have shown small chemical changes in their AG-binding pocket perturb susceptibility to AGs. Methylation at N7-G1405 in *E. coli*, and at analogous nt residues in *Klebsiella pneumoniae, Pseudomonas aeruginosa* and *Streptomyces tenebrarius* (G1388 in *M. smegmatis*) induces resistance to 4,6-disubstituted and 4,5-disubstituted AGs (34-36). Also, methylation at C1407 (C1390 in *M. smegmatis*) and A1408 (A1391 in *M. smegmatis*) cause kanamycin and gentamicin resistance in pathogenic *E. coli* (37, 38). Hence, the difference in kanamycin (and related AGs) sensitivity observed between the wild-type and ΔbS22 strains of *M. smegmatis* can be attributed to the observed change in conformations of nts 1388-1392 in h44.

Despite the extensive conservation of both small ribosomal proteins, bS22 and bL37, a defined role for either protein has yet to be determined. In this study, we focus on the mycobacterial protein, bS22, and find that is not essential for cell viability, similar to its eukaryotic and mitochondrial orthologs, eL41 and mS38, respectively. We find that loss of bS22 confers increased sensitivity to AG antibiotics kanamycin, apramycin, gentamycin and streptomycin. To address the mechanism behind this observation, we report the first structure of kanamycin in a complex with a mycobacterial ribosome. The solved structures and AG susceptibility profiles are consistent with local conformational changes of h44 nts that result from bS22 binding also impair kanamycin interactions. This displacement of h44 is analogous to the roles played by substitutions of Lys42 and Lys87 in *E. coli* uS12, which alter streptomycin binding and confer streptomycin resistance (39).

## Materials and Methods

### Ribosome purification

*M. smegmatis* wild-type and ΔbS22 ribosomes were purified as described previously (40). Cells harvested from 500 ml cultures of *M. smegmatis* strains were flash-frozen in liquid nitrogen. A miller (Retsch MM400) was used to pulverize the frozen cells for 6 cycles. In each cycle, 3 min of milling at 15 Hz was followed by 30 s of cooling in liquid nitrogen. The pulverized cells were resuspended in a low-salt buffer, HMA-10 (20 mM HEPES-K [pH 7.5], 30 mM NH4Cl, 10 mM MgCl2, 5 mM β-mercaptoethanol). The lysate was centrifuged for 30 min at 30,000 × g in a Thermo Sorvall Lynx 4000 centrifuge at 4°C. The supernatant was collected in ultracentrifuge tubes (Beckman PC) and centrifuged at 42,800 rpm, 4°C for 2 h 15 min in a Beckman rotor (type 70Ti). The pellet obtained was soaked over-night in 4 ml of low salt HMA-10 buffer and then homogenized in a pre-chilled glass homogenizer. 3 units/ml of RNase-free Turbo DNase (Invitrogen) was added to the homogenate and incubated for 1 h at 4°C. Thereafter, 4 ml of high-salt HMA-10 buffer (20 mM HEPES-K [pH 7.5], 600 mM NH4Cl, 10 mM MgCl2, 5 mM β-mercaptoethanol) was added followed by incubation at 4°C for 2 h. The mixture was centrifuged at 20,000 × g for 15 min in a Thermo Sorvall Lynx 4000 centrifuge. The supernatant was centrifuged for 2 h 15 min at 42, 800 rpm in a Beckman rotor type 70Ti at 4°C to collect the crude ribosome pellet. The pellet was resuspended in low-salt HMA-1 (20 mM HEPES-K [pH 7.5], 30 mM NH4Cl, 1 mM MgCl2, 5 mM β-mercaptoethanol) buffer, and layered on a 40-ml sucrose gradient (10% to 40%) prepared in the same buffer and centrifuged for 16 h at 24,000 rpm in a Beckman rotor SW28. The density gradient fractionation system (Brandel) was used for fractionation and the fractions corresponding to 30S were pooled. The pooled 30S fractions were pelleted by ultracentrifugation at 42,800 rpm for 4 h in Beckman rotor Type 70Ti and resuspended in storage buffer.

### Construction of gene deletions and *bs22* mutations

Deletions of genes MSMEG_0945 (*bL*22 and MSMEG_1916 (*bL37*) were creatd by recombineering (22). Substrates for recombineering were made by Soeing-PCR that combined ∼ 500 bp of 5’ and 3’ flanking DNA with a cassette encoding zeocin resistance (*zeo*), which was designed to precisely replace the target gene (23). Recombinants were confirmed by PCR using flanking primers.. The *zeo* gene was subsequently excised from the chromosome by Cre-mediated site-specific recombination between two *loxP* sites included at the 5’ and 3’ ends of the *zeo* cassette. The modified loci were confirmed by Sanger sequencing of PCR amplification products spanning the targeted genes. Thus, the two strains used to assess the functions of bL37 and bs22 contain a single *loxP* sitein place of each gene. The strain with replacement of both genes was derived from the *ΔbS22* knock out. MSMEG_0945 (*bs22*) and MSMEG_1916 (*bl37*) were cloned for complementation and mutagenesis studies by PCR into the E. coli-mycobacterial shuttle vector pGD6, which encodes resistance to hygromycin (41). Each clone was confirmed by Sanger sequencing. Oligonucleotide sequences used for cloning are available on request

### Phenotypic assays

Growth competition assays used zeo-marked deletion strains and a hygromycin-marked, wild-type strain (MKD158; (42)). MKD158 and mutant strains were grown to saturation overnight, diluted and combined, such that each contributed equal OD600 units to the starting culture. 10 ml cultures of Trypticase Soy broth with Tween-80 (TSBT) were shaken in a 50 ml flask and passaged by sub-culturing 1:1,000 once a day for four days. An aliquot of cells was taken at T=0 and at each subculture time point, diluted, and plated on antibiotic medium selecting for either mutant (zeocin resistant) or wild-type (hygromycin resistant) cells. Colonies were enumerated and relative survival plotted. These assays were performed in duplicate at 30^°^, 37^°^and 45^°^C. Assays were also done in minimal medium (Sauton’s broth + TPEN) to chelate zinc, which is known to impact mycobacterial ribosomal protein content (18). Cells were also grown with sub-inhibitory concentrations of spectinomycin (6.25 *μ*g/ml) and erythromycin (0.156 *μ*g/ml) to exert pharmacologic stress on the ribosome. None of the growth conditions resulted in an obvious outgrowth of wild-type over mutant cells.

### Biofilm assays

Biofilm assays were performed in 6-well dishes using either TSB or complete biofilm medium with 1 *μ*M TPEN to remove zinc and with 1 mM zinc sulphate (43). A 10 *μ*l aliquot of an overnight culture was used to inoculate 5 ml of medium in each well. Biofilms were incubated for up to 10 days with minimal handling at 30^°^C and evaluated. The mutant strains developed and formed biofilms in the same timeframe as the wild-type strain and with the same overall architecture, regardless of medium.

### Disk diffusion assays

Wild-type and *Δbs22* mutant strains were grown to an OD600 of 0.6 in TSBT. 200 *μ*l of this culture was added to 3 ml of top agar and poured onto a TSA plate without antibiotic. The medium was allowed to set and stored at 4^°^C. Paper disks were placed on the ungrown lawn and 5 *μ*l of test reagents were soaked into the disk. Plates were incubated at 37^°^C overnight and the zone of inhibition scored relative to wild-type cells. The concentrations of antibiotics tested were: azithromycin 2 *μ*g/ml, apramycin 0.25 *μ*g/ml, chloramphenicol 20 *μ*g/ml, erythromycin 15 *μ*g/ml, gentamycin 0.2 *μ*g/ml, kanamycin 0.25 *μ*g/ml, retapamulin 10 *μ*g/ml, streptomycin 0.24 *μ*g/ml, tetracycline 1 *μ*g/ml, and zeocin 1 *μ*g/ml.

### MIC assays

Wild-type and *Δbs22* strains were grown to an OD600 of 0.6 in TBST. Cultures were diluted to an OD of 0.08 and 50 *μ*l used to inoculate 2 ml of TBST containing 2-fold serial dilutions of the antibiotic to be tested. The cells were incubated at 37 ^°^C with shaking for 36 hrs and the MIC determined by culture growth compared to the wild type at matched antibiotic concentrations.

### RNA-seq and Ribo-seq

This was performed on the *ΔbS22* and wild-type strains of *M. smegmatis* using the protocols described in (24, 25, 44). Briefly, overnight cultures of each strain were used to inoculate 400 mls of 7H9 medium with Tween-80 and 0.2% glycerol. Cells were grown shaking to an OD_600_ of 1.0. Cells were harvested by rapid filtration using a 0.45 mm PES filter system (Cell treat) and flash frozen in liquid nitrogen in lysis buffer (44). Frozen cells were pulverized 6 times at 15 Hz for 3 min in a mixer mill (Retsch MM4000). An aliquot of the pulverized cells was kept for RNA-seq. For RNA-seq, 16S and 28S ribosomal RNAs were removed by subtractive hybridization from this aliquot before fragmenting the RNAs and converting into cDNA libraries for sequencing. The remaining material was treated with micrococcal nuclease to degrade DNA and monosomes were isolated by fractionation on a sucrose gradient. The monosome fraction was treated with phenol-chlorofom to remove protein and RNA collected by isopropanol precipitation. RNA was then converted into cDNA, PCR amplified with appropriate adapters and deep sequenced using lllumina HiSEq technology(44, 45). Raw reads for both RNA- and Ribo-seq were processed as described previously (24, 25).

### Cryo-electron microscopy and image processing

4 μl of 30S wild-type or ΔbS22 ribosomal subunits (150 nM), in the presence or absence of kanamycin (2 mM) was applied to Quantifoil holey copper 1.2/1.3 grids, pre-coated with a thin layer (∼50 Å thick) of continuous carbon film, and glow-discharged for 30 s on a plasma sterilizer. The sample was incubated on the grids for 15 s at 4 °C and 100% humidity, then blotted for 4 s and immediately plunged into liquid ethane for vitrification with the help of a Vitrobot (FEI company). Data were collected on a Titan Krios electron microscope (FEI company) equipped with a Gatan K2 summit direct-electron detecting camera at 300 kV. Data collection parameters for each sample are listed in Supplementary Table 1. CryoSPARC pipeline (46) was used for data processing. Full-frame motion correction was applied to all movie frames corresponding to the micrographs. CTFFIND464 was used to determine the contrast transfer function (CTF) and bad images were eliminated at this step. The remaining micrographs were used for particle picking using the auto-pick function. Local motion correction was performed and was followed by reference-free 2D classification. Appropriate 2D-class averages were selected and processed further. Ab initio reconstructions and reference-based 3D classifications were done subsequently (as specified in the data-processing pipelines **Fig. S2-4**) to resolve particles into the 70S, 50S, and 30S (**Fig. S4**) and also to obtain homogenous sub-populations of 30S (**Fig. S2-3**). Finally, 30S classes containing 80,416 particles from the ΔS22-30S-kanamycin dataset, 179,811 particles from the ΔS22-30S dataset, and 70,961 particles from the wild-type 30S dataset were refined to 3.1 Å, 3.1 Å, and 3.8 Å, respectively.

### Model building and optimization

Coordinates corresponding to the small ribosomal subunit of the *M. smegmatis* ribosome (PDB ID: 6DZK, 5O5J) was used as the initial template. The higher resolution of our maps (**Fig. S2-S4, S5**) enabled us to build most rRNA and ribosomal proteins. The initial template was modified and fitted manually into the respective maps using UCSF Chimera 1.14 and COOT. Subsequently the models were real-space refined in COOT (47). This was followed by refinement using “phenix.real_space_refine” (48). The models were validated using PHENIX “Comprehensive validation”. The structural graphics are generated using UCSF-Chimera and Chimera-X and arranged in Adobe Illustrator.

## Acknowledgements

This work was supported by grants to RKA (NIH:GM61576) and jointly to KMD, TAG, and JTW (NIH: GM GM139277). RKA. also acknowledges support to his lab through NIH R01 grants AI132422, GM139277 and AI155473. Support from the Wadsworth Center core facilities–Applied Genomic Technologies and the media core facility– is acknowledged. We also acknowledge Wadsworth Center’s and New York Structural Biology Center’s (NYSBC’s) 3D-EM facilities. NYSBC EM facilities are supported by grants from the Simons Foundation (349247), NYSTAR, the NIH (GM103310) and the Agouron Institute (F00316).

## Authors’ Contribution

AD, MRS, RKK, and RKA designed the structural studies. JTW, TAG, and KMD designed the biochemical and genetics studies. SM, AD, MRS, RKK, PK, NKB, and RKA participated cryo-EM structure determination. SM, MRS, and RKA performed the structural analysis. JC, MS, CM, PK, JTW, TAG, and KMD performed and interpreted the genetics and biochemical experiments; SM, JTW, TAG, KMD and RKA wrote the manuscript, and all other authors read and commented on the manuscript.

## Data Deposition

The cryo-EM maps and atomic coordinates of the kanamycin-bound and unbound delta bS22 30S subunits are deposited in the Electron Microscopy and PDB Data Bank (wwPDB.org). Accession codes for kanamycin-bound and unbound delta bS22 30S subunit maps are EMD-29457 and EMD-29480, respectively. Accession codes for coordinates of kanamycin-bound and unbound delta bS22 30S subunit are PDB ID 8FUE and 8FUS, respectively.

