## Supplementary Figures and a Table for "The small mycobacterial ribosomal protein, bS22, modulates aminoglycoside accessibility to its 16S rRNA helix-44 binding site"

Divisions of [1]Translational Medicine and [2]Genetics, Wadsworth Center, New York State Department of Health, Albany, NY 12237
and
[3]Department of Biomedical Sciences, University at Albany, SUNY, Albany, NY 12222.

*equal contributors
[#] corresponding author

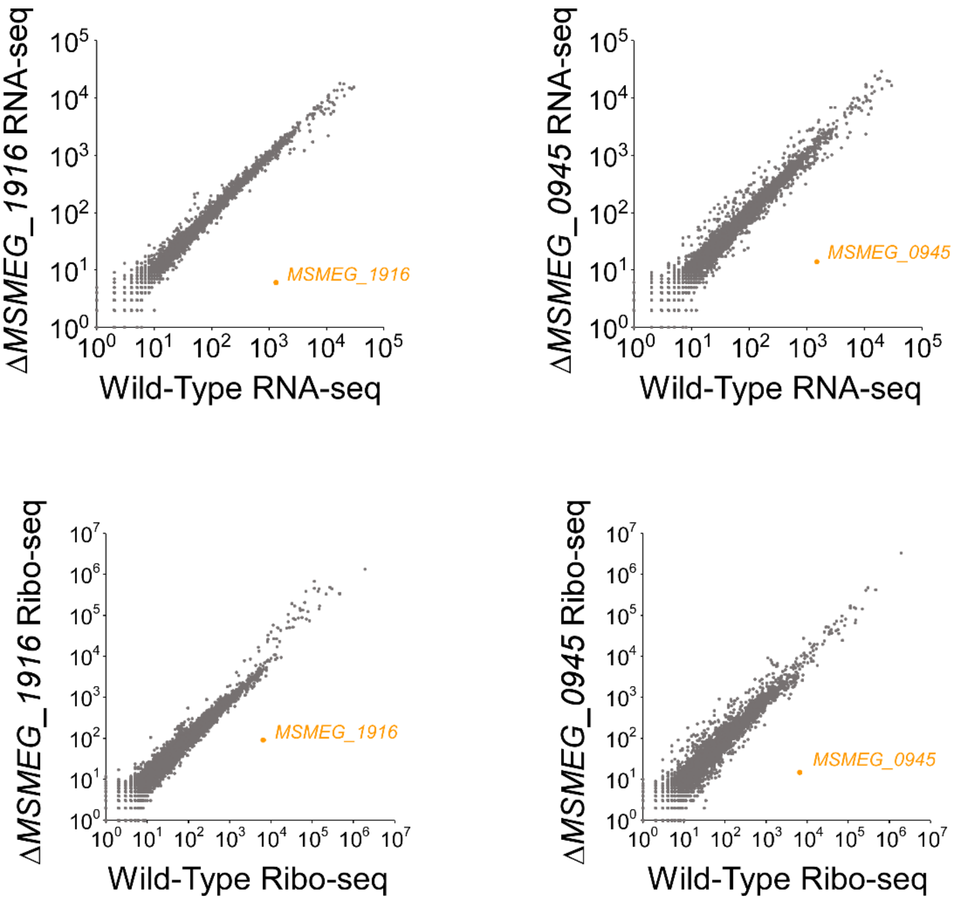


**Figure S1. RNA-seq and Ribo-seq analysis of *M. smegmatis* ΔMSMEG_1916 and *M. smegmatis* ΔMSMEG_0945 relative to wild-type.** Scatter-plots show RNA levels (RNA-seq) or ribosome-associated RNA levels (Ribo-seq) for all M. smegmatis genes, comparing between wild-type and ΔMSMEG_1916, and between wild-type and ΔMSMEG_0945. Genes are represented by gray circles, with the deleted gene represented by an orange circle.

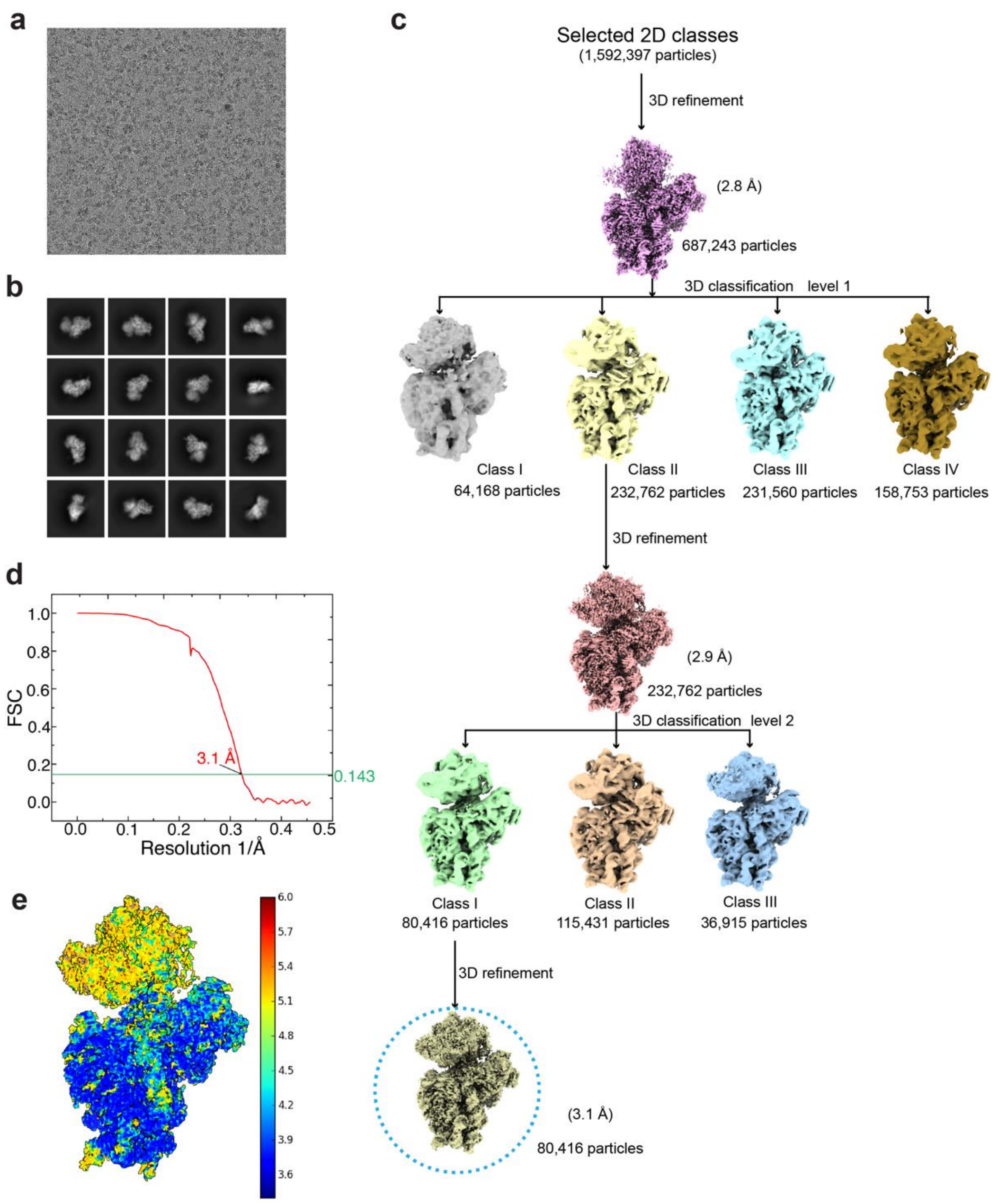


**Figure S2. The cryo-EM image reconstruction pipeline for the *M. smegmatis* ΔS22-30S-kanamycin complex.** (a) A representative micrograph from the *M. smegmatis* ΔS22-30S-kanamycin dataset. (b) Representative 2D class averages selected for the 3D reconstructions. (c) Flowchart depicting the 3D classifications and refinements. The final map (80,416 particles) used in this study is circled (blue dashes) (d) Fourier-shell correlation (FSC) plot for the final map (circled in blue dashes in panel c). (e) Local resolution of the final map of *M. smegmatis* ΔS22-30S-kanamycin complex.

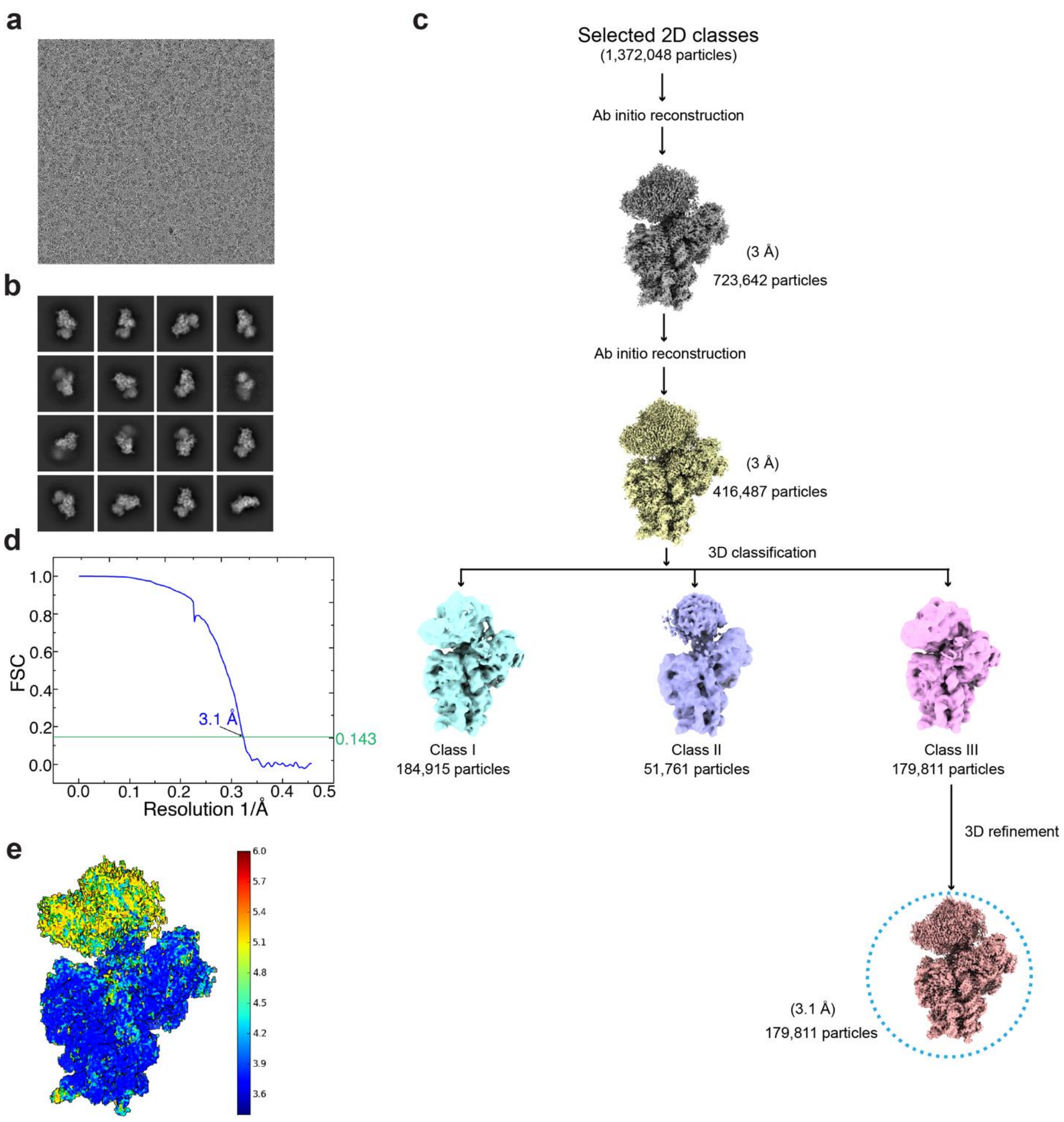


**Figure S3. The cryo-EM image reconstruction pipeline for *M. smegmatis* ΔS22-30S.** (a) A representative micrograph from the *M. smegmatis* ΔS22-30S dataset. (b) Representative 2D class averages selected for the 3D reconstructions. (c) Flowchart depicting the 3D classifications and refinements. The final map (179,811 particles) used in this study is circled (blue dashes) (d) Fourier-shell correlation (FSC) plot for the final map (circled in blue dashes in panel c). (e) Local resolution of the final map of *M. smegmatis* ΔS22-30S.

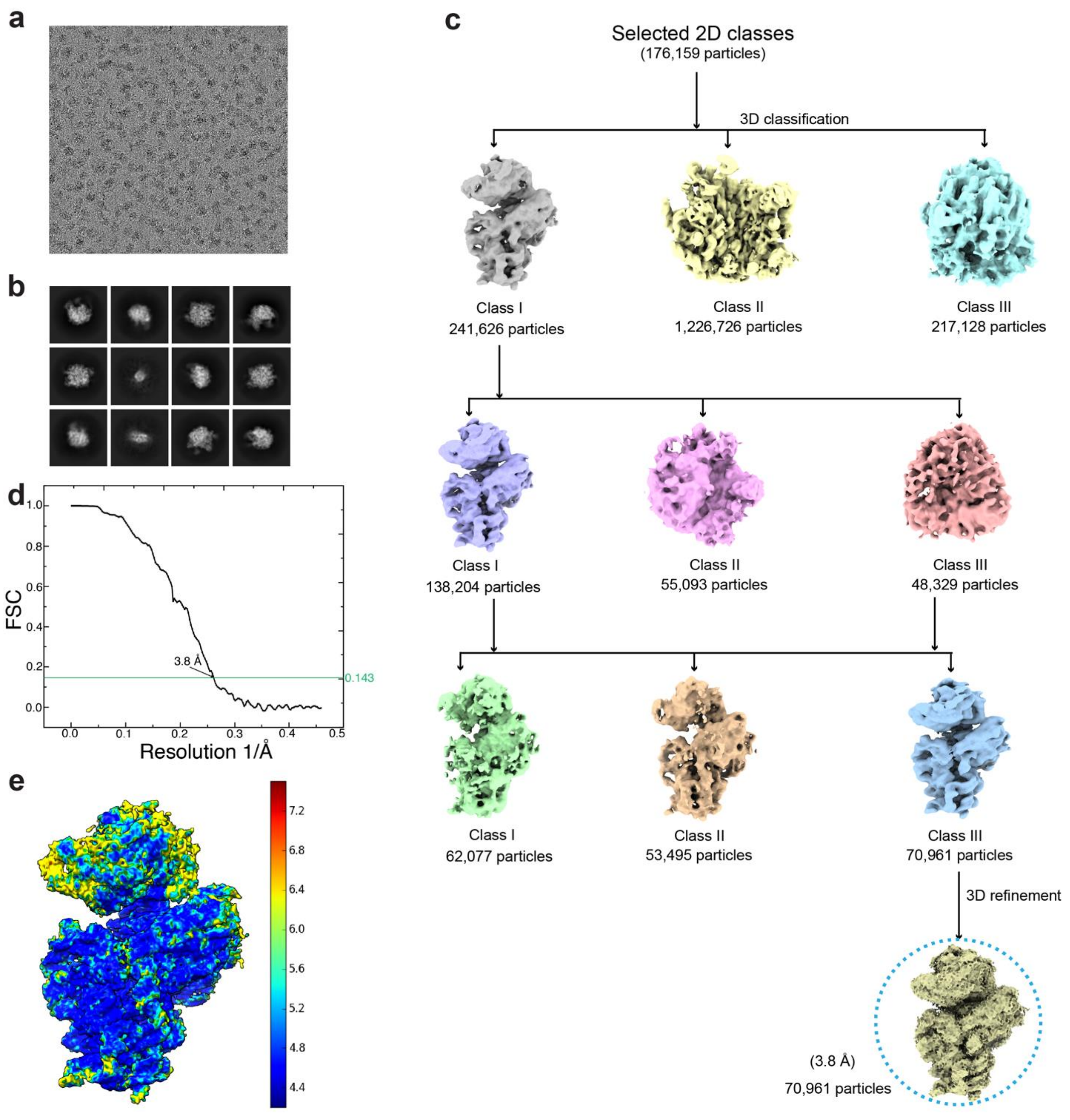


**Figure S4. The cryo-EM image reconstruction pipeline for wild-type 30S.** (a) A representative micrograph from the *M. smegmatis* wild-type ribosome dataset. (b) Representative 2D class averages selected for the 3D reconstructions. (c) Flowchart depicting the 3D classifications and refinements. The final map (70,961 particles) used in this study is circled (blue dashes) (d) Fourier-shell correlation (FSC) plot for the final map (circled in blue dashes in panel c). (e) Local resolution of the final map of *M. smegmatis* wild-type 30S.

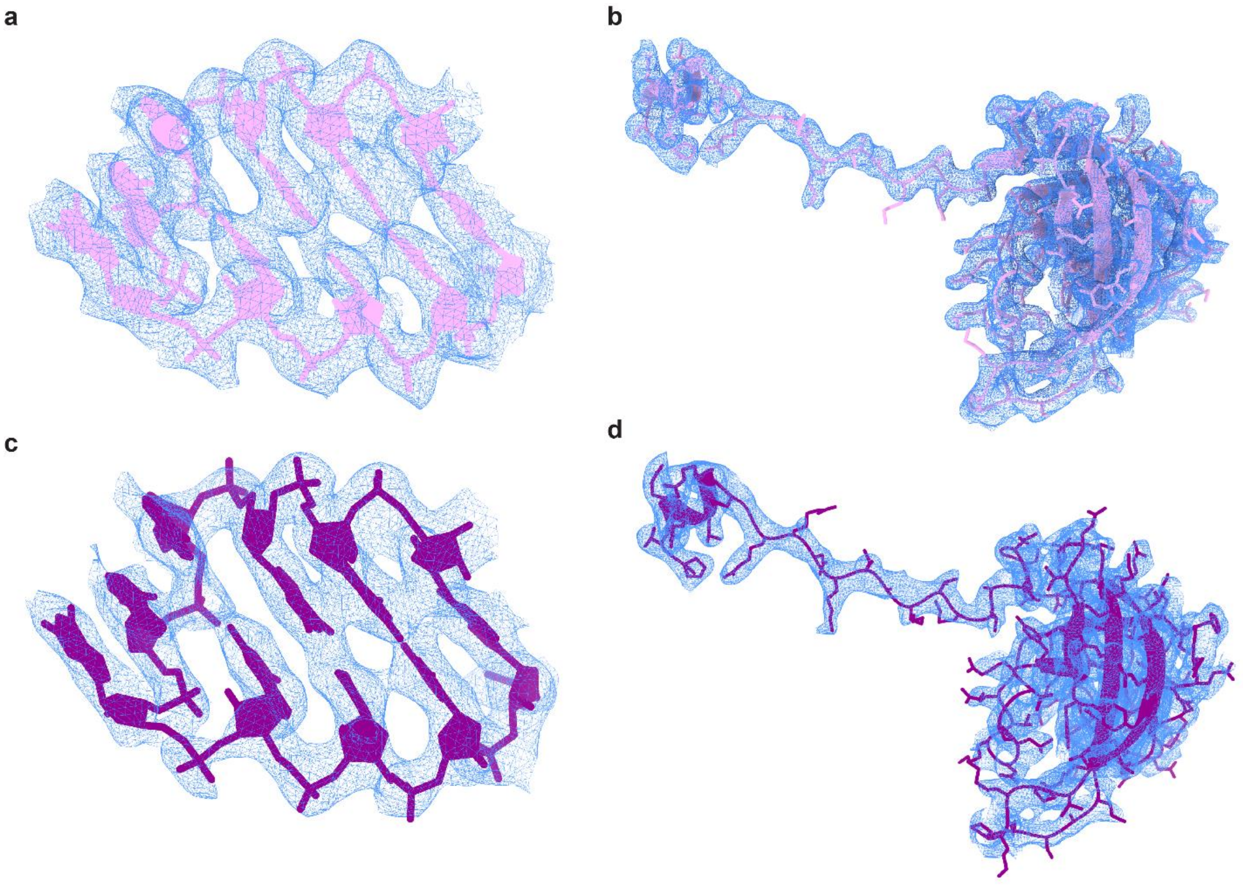


**Figure S5. Representative segments from the cryo-EM density maps display high-resolution features.** Cryo-EM density and model corresponding to h45 of 16S rRNA from (**a**) ΔS22 30S-kanamycin complex and (**c**) ΔS22 30S. Cryo-EM density and model corresponding to uS12 from (**b**) ΔS22 30S-kanamycin complex and (**d**) ΔS22 30S.

**Table S1. Data collection, Refinement and Model Validation parameters.**

| **Description** | **ΔS22-30S-Kan complex** | **ΔS22-30S** |
|---|---|---|
| **Data collection** | | |
| Microscope | FEI Titan Krios | FEI Titan Krios |
| Voltage (kV) | 300 | 300 |
| Pixel size (Å) | 1.083 | 1.083 |
| Defocus range (μm) | 0.8-2.5 | 1.0-2.5 |
| Average $e^-$ dose per image ($e^-$/ Å$^2$) | 50.92 | 51.27 |
| Particles (initial) | 3,201,547 | 2,244,260 |
| Particles (final) | 80,416 | 179,811 |
| FSC-threshold | 0.143 | 0.143 |
| Resolution (Å) | 3.1 | 3.1 |
| Map-sharpening B factor (Å$^2$) overall | 59.9 | 93.4 |
| **Refinement** | | |
| RMS deviations | | |
| Bonds (Å) | 0.007 | 0.005 |
| Angles ($^o$) | 0.744 | 0.652 |
| MolProbity score | 2.17 | 2.10 |
| Clash score | 12.45 | 11.41 |
| Rotamer outliers (%) | 0.56 | 0.61 |
| Ramachandran plot | | |
| Outliers (%) | 0.09 | 0.00 |
| Allowed (%) | 10.54 | 9.61 |
| Favored (%) | 89.38 | 90.84 |
| RNA | | |
| Correct sugar puckers (%) | 99.07 | 98.94 |
| Angle outliers (%) | 0.00 | 0.00 |
| Bond outliers (%) | 0.00 | 0.00 |
| Good backbone conformations (%) | 75.84 | 76.51 |
| Model composition | | |
| RNA bases | 1,511 | 1,511 |
| Protein residues | 2,363 | 2,363 |
